# Lights, Camera, Path Splitter: A New Approach for Truly Simultaneous Dual Optical Mapping of the Heart with a Single Camera

**DOI:** 10.1101/651380

**Authors:** Rafael Jaimes, Damon McCullough, Bryan Siegel, Luther Swift, James Hiebert, Daniel McInerney, Nikki Gillum Posnack

## Abstract

**Background:** Optical mapping of transmembrane voltage and intracellular calcium is a powerful tool for investigating cardiac physiology and pathophysiology. However, simultaneous dual mapping of two fluorescent probes remains technically challenging. We introduce a novel, easy-to-use approach that requires a path splitter, single camera and excitation light to simultaneously acquire voltage and calcium signals from whole heart preparations, which can be applied to other physiological models – including neurons and isolated cardiomyocytes.

**Results:** Complementary probes were selected that could be excited with a single wavelength light source. Langendorff-perfused hearts (rat, swine) were stained and imaged using a sCMOS camera outfitted with an optical path splitter to simultaneously acquire two emission fields at high spatial and temporal resolution. Voltage (RH237) and calcium (Rhod2) signals were acquired concurrently on a single sensor, resulting in two 384×256 images at 814 frames per second. At this frame rate, the signal-to-noise ratio was 47 (RH237) and 85 (Rhod2). Imaging experiments were performed on small rodent hearts, as well as larger pig hearts with sufficient optical signals. In separate experiments, each dye was used independently to assess crosstalk and demonstrate signal specificity. Additionally, the effect of ryanodine on myocardial calcium transients was validated – with no measurable effect on the amplitude of optical action potentials. To demonstrate spatial resolution, ventricular tachycardia was induced –resulting in the novel finding that spatially discordant calcium alternans can be present in different regions of the heart, even when electrical alternans remain concordant. The described system excels in providing a wide field of view and high spatiotemporal resolution for a variety of cardiac preparations.

**Conclusions:** We report the first multiparametric mapping system that simultaneously acquires calcium and voltage signals from cardiac preparations, using a path splitter, single camera and excitation light. This approach eliminates the need for multiple cameras, excitation light patterning or frame interleaving. These features can aid in the adoption of dual mapping technology by the broader cardiovascular research community, and decrease the barrier of entry into panoramic heart imaging, as it reduces the number of required cameras.

## BACKGROUND

Cardiovascular research has been propelled by the advent of parameter-sensitive probes, which can be used to monitor transmembrane voltage and intracellular calcium within live cardiac preparations (1–5). Optical mapping is an imaging technique that measures fluorescence signals across a cardiac preparation with high spatiotemporal resolution. Optical mapping of voltage-sensitive probes (6–8) allows for the measurement of action potential morphology and the spread of electrical activity, as well as the identification of tissue heterogeneities that can promote arrhythmias. Whereas intracellular calcium probes (9) are used to investigate modifications in excitation-contraction coupling, which can alter action potential duration, elicit after-depolarizations, and promote electrical/mechanical alternans. Accordingly, simultaneous imaging of both transmembrane voltage and intracellular calcium is a powerful integrative tool for cardiac research (for extensive reviews see(1–4)). Yet, assembling a dual optical mapping system remains technically challenging (4,10,11), which has limited the use of this technique to a relatively small number of research laboratories.

### Dual camera configurations

Simultaneous dual mapping systems have traditionally used a dual-sensor design, wherein the emission of each complementary probe (voltage, calcium) is separated by wavelength and diverted to two separate detectors (1,5,12–16). Such a design was described by Fast and Ideker, in which cardiomyocytes monolayers were stained with complementary probes (RH237 – voltage, Fluo-3AM – calcium) and the emitted fluorescent signals were focused onto two 16×16 photodiode arrays(15). A similar approach was employed by Choi and Salama to simultaneously record transmembrane voltage (RH237) and intracellular calcium (Rhod2) signals from isolated, whole guinea pig hearts (13). Subsequently, dual-sensor optical configurations have been expanded to include EMCCD and sCMOS sensors with improved spatial resolution (see Table 1 for example configurations (4,5,17)). RH237/Rhod2-AM probes are still commonly used for dual imaging (1,18–22), although dual-dye combinations that separate fluorescence signals by emission have also been developed. They include: Di-4-ANEPPS with Indo-1(16), Di-2-ANEPEQ and Calcium green (23), and RH237 with Fluo-3/4/5N (12,15,24,25). Importantly, these dye combinations can have spectral overlap, which necessitates non-ideal emission bandpass to negate spectral overlap and/or the inclusion of a calcium probe with an inferior dissociation constant(26). A dual-sensor optical mapping system offers many advantages, including the full spatial and temporal resolution of each individual camera. However, a dual-camera optical setup can be both technically challenging and cost-prohibitive for basic-science and teaching laboratories (see Table 1 (3,4,10,11)). Dual-sensor systems also require proper alignment to ensure that fluorescence signals are being analyzed from the same tissue region on each individual detector, which could lead to erroneous results. Finally, the physical footprint of two cameras in a 90 degree orientation can be limiting and reduce its versatility.

**Table 1:**
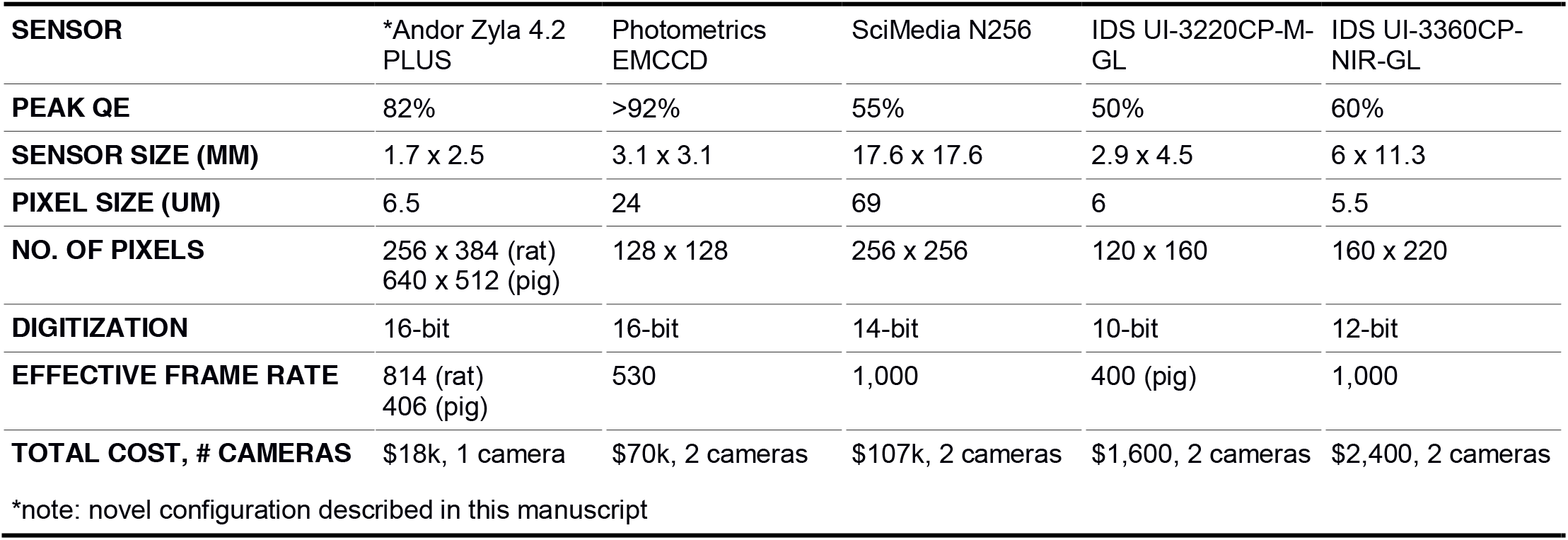
Example of optical mapping configurations using dual sensors

| SENSOR | *Andor Zyla 4.2 PLUS | Photometrics EMCCD | SciMedia N256 | IDS UI-3220CP-M-GL | IDS UI-3360CP-NIR-GL |
| --- | --- | --- | --- | --- | --- |
| PEAK QE | 82% | >92% | 55% | 50% | 60% |
| SENSOR SIZE (MM) | 1.7 x 2.5 | 3.1 x 3.1 | 17.6 x 17.6 | 2.9 x 4.5 | 6 x 11.3 |
| PIXEL SIZE (UM) | 6.5 | 24 | 69 | 6 | 5.5 |
| NO. OF PIXELS | 256 x 384 (rat)<br>640 x 512 (pig) | 128 x 128 | 256 x 256 | 120 x 160 | 160 x 220 |
| DIGITIZATION | 16-bit | 16-bit | 14-bit | 10-bit | 12-bit |
| EFFECTIVE FRAME RATE | 814 (rat)<br>406 (pig) | 530 | 1,000 | 400 (pig) | 1,000 |
| TOTAL COST, # CAMERAS | \$18k, 1 camera | \$70k, 2 cameras | \$107k, 2 cameras | \$1,600, 2 cameras | \$2,400, 2 cameras |

\*note: novel configuration described in this manuscript

### Single camera configurations

An alternative optical mapping approach includes the use of a single sensor to *sequentially* image each complementary probe (voltage, calcium) in time using excitation light patterning (10,11,27,28) (see Table 2). A single-sensor, sequential imaging approach was described by Lee et al., which achieved rapid excitation light switching by utilizing recently developed high-power light emitting diodes (10). Accordingly, single-sensor designs use dual-dye combinations that require two (or more) excitation light sources, but share a single emission band, including: di-4-ANBDQPQ and Fura-2 (10) or Rhod2-AM (29), Di-4-ANBDQBS and Fluo-4 (28), or Di-4-ANEPPS and X-Rhod-1(11) (for review see (26)). Single-sensor optical mapping systems offer a cost advantage, since the camera sensor is often one of the most expensive components of an optical mapping setup. However, a single-sensor platform design is technically challenging since the use of two different excitation light wavelengths necessitates light source triggering, camera synchronization and frame interleaving (10,27,28). A microcontroller is typically used to properly time light triggering and to coordinate image acquisition, since each (alternating) probe signal is acquired in sequential frames. This alternating configuration decreases the frame rate for each fluorescent probe of interest, since voltage/calcium signals typically reside on odd/even images. Excitation light ramp up/down times and shutter open/close times also have to be taken into consideration to avoid overlap. Accordingly, single-sensor setups (with interleaved frames per dye of interest) necessitate shortened exposure times compared with dual-sensor setups. The latter can diminish signal-to-noise quality, without offering the same temporal fidelity.

**Table 2:**
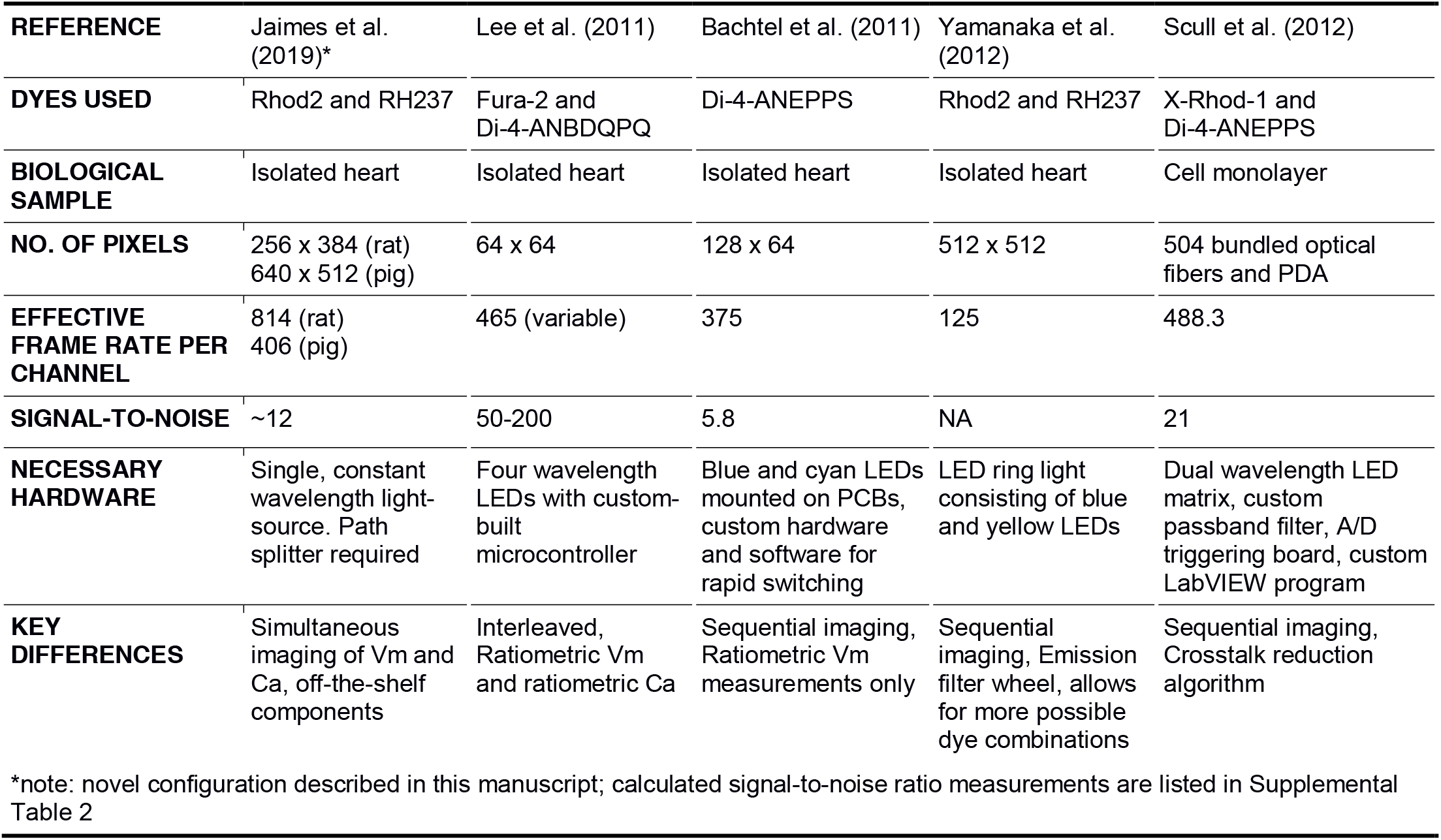
Optical mapping experiments employing a single sensor design.

In the described study, we describe the first multiparametric mapping system that simultaneously acquires calcium and voltage signals from cardiac preparations, using a commercially available optical path splitter, single camera and single excitation light (Figure 1,2). Specifically, we have taken advantage of recently developed large field of view sCMOS sensors that are faster and more sensitive (2048 × 2048 pixels, 18.8 mm diagonal, Zyla 4.2 plus, Andor Technology PLC, Belfast, UK). Our configuration separates the two emission bands for Rhod2 and RH237, however, we negate the need for a bulkier footprint and costly two camera setup by simultaneously directing each emission band to different sides of the single, large camera sensor. To date, optical path splitters have largely been limited to microscopy applications that allow for a slow speed of acquisition (longer exposure time) and utilize bright immunofluorescent samples. However, such an approach has not yet been described for multiparametric imaging of whole heart preparations that require 1) fast acquisition speeds, 2) high spatial resolution, and 3) utilize fluorescent probes with low signal to noise ratios.

**Figure 1:**
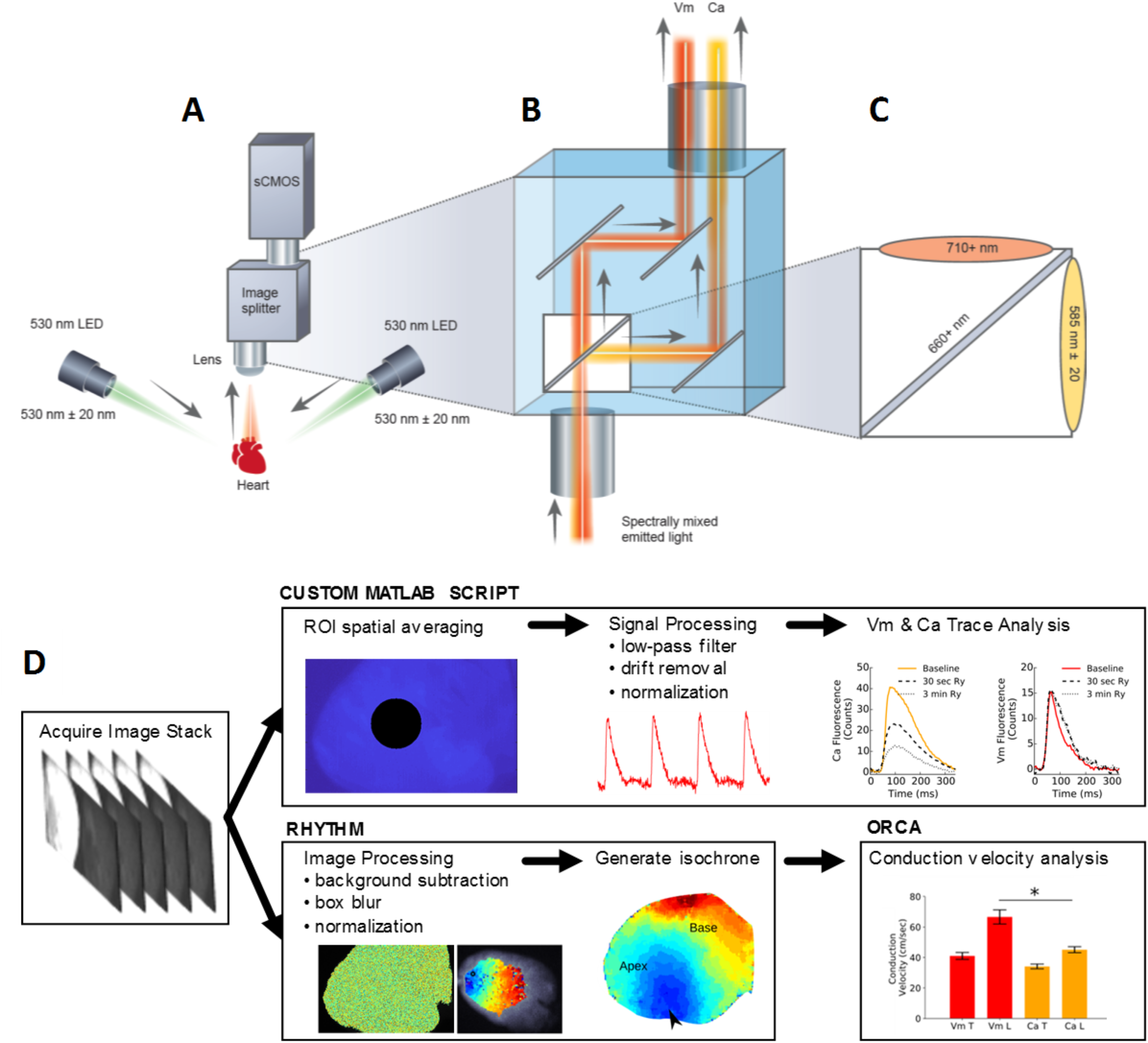
Overview of the optical design and workflow. (**A**) Optical system configuration with image splitting device positioned in front of a sCMOS camera (Zyla 4.2 PLUS, Andor Technologies), (**B**) Emission of each complementary probe (Vm, Ca) is separated by wavelength using an image splitting device (**C**) Dichroic cube setup with the two emission filters and a dichroic mirror. (**D**) Experimental workflow includes image acquisition, followed by signal processing using a custom MATLAB script or image processing in Rhythm(38) and conduction velocity analysis in ORCA(58). Vm = transmembrane voltage, Ca = intracellular calcium signal.

**Figure 2:**
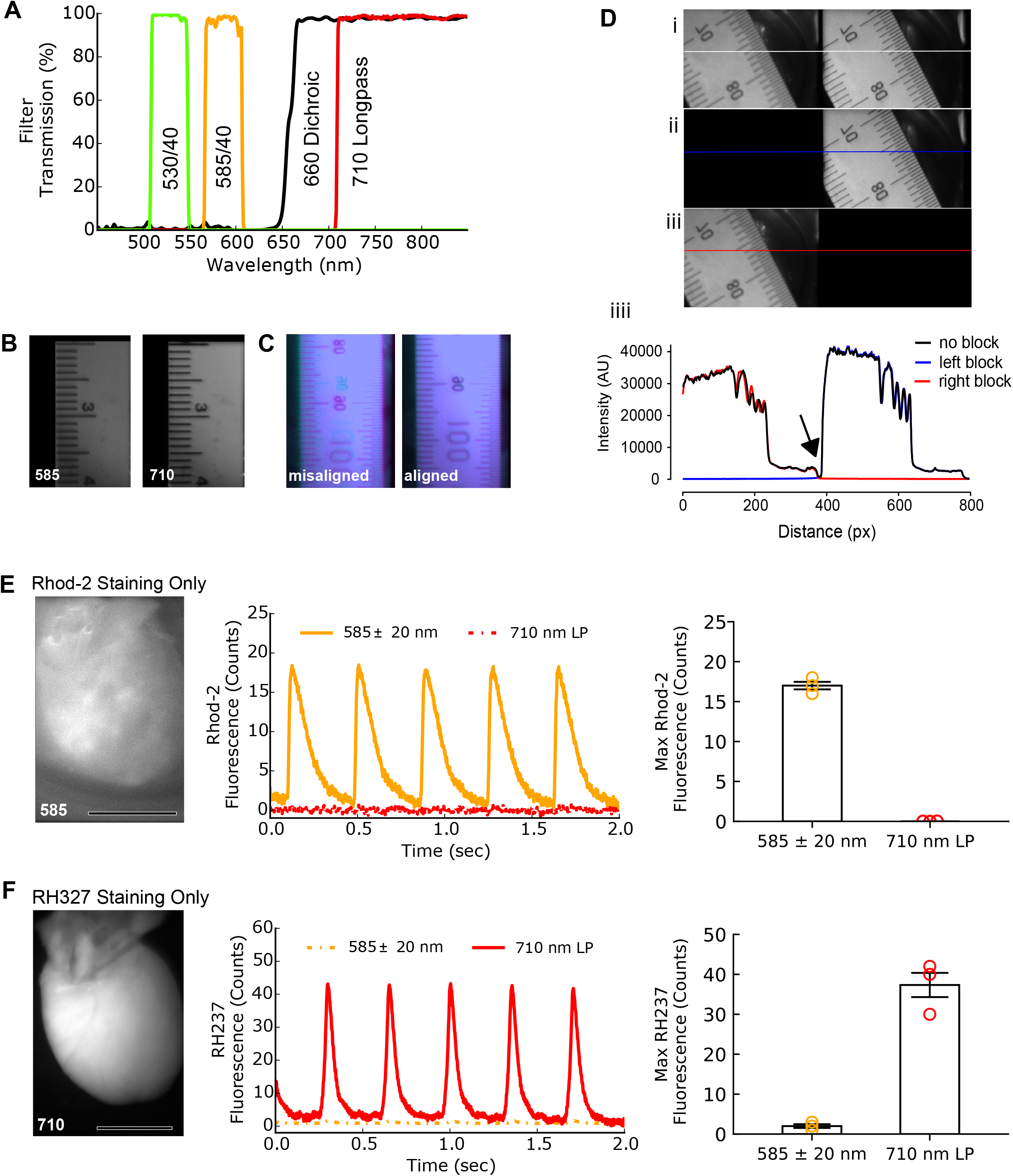
Optical configuration results in negligible crosstalk between fluorescent probes. (**A**) Passbands of all the filters used in the optical configuration. (**B**) Two sides of a single, large camera sensor display a duplicated ruler image from each emission band with negligible focal shift. (**C**) Illustration of misaligned (right) and proper alignment (left) of images. (**D**) **i**: 800×256 pixel image of a back illuminated ruler is shown before splitting into two 384×256 pixel images; left: long wavelength path (710+nm), middle: black boundary between images (16 pixels), right: short wavelength path (585±20nm). **(ii)** 800×256 image when the a plate is inserted to block the light path on the left (710+nm). **(iii)** 800×256 image when the path on the right is blocked (585 ± 20nm). **(iiii)** Profile traces corresponding to the full image (black), left path blocked (blue, 710+nm) and right path blocked (red, 585±20nm) demonstrate a lack of crosstalk along the light paths. Arrow denotes location of the boundary between images without crosstalk. (**E**) Left: Detected fluorescence on the epicardial surface after single probe administration (Rhod2). Center: High quality calcium transients through the 585 nm emission filter with no detectable signal through the 710 nm longpass filter. Right: Maximum counts from each channel after Rhod2 (only) administration. (**F**) Left: Detected fluorescence on the epicardial surface after single probe administration (RH237). Center: High quality optical action potentials through the 710 nm emission filter, with minimal signal through the 585 nm emission. Right: Maximum counts from each channel after RH237 (only) administration. Note: Bar plots (E,F) on the right collected from different rat hearts, each loaded with a single fluorescent probe (n = 3 each plot). Scale bar = 1cm.

Notably, the described approach enables truly simultaneous dual imaging of cardiac preparations, while eliminating the need for two cameras and/or multiple excitation light sources, light patterning and frame interleaving. The described imaging platform is composed entirely of off-the-shelf components, which can aid in the adoption and successful assembly of this setup by other laboratories. The described protocol also employs a commonly used dual-dye combination (RH237, Rhod2-AM) that is widely available, thereby negating the need for genetically-encoded indicators (30–32) or fluorescent probes that are not yet commercially available (33,34). We validate the utility of our approach by performing high-speed simultaneous dual imaging with sufficient signal-to-noise ratio for calcium and voltage signals and specificity of emission signals with negligible cross-talk. Ventricular tachycardia was induced to demonstrate high spatiotemporal resolution, which resulted in a novel finding that spatially discordant calcium alternans can be present in different regions of the heart, even when electrical alternans remain concordant. Furthermore, we highlight the versatility of our imaging platform by seamlessly maneuvering our optical setup between a Langendorff-perfused rat heart setup (2-3 cm in length, laying down) and Langendorff-perfused piglet heart setup (5-8 cm in length, suspended) with different orientations. Due to technical challenges, dual optical mapping of a larger pig heart has only been previously described by one other group (35); albeit this study employed a traditional dual camera configuration with limited spatial resolution (see Table 1). Taken together, the described dual imaging platform may be of interest to a wide variety of basic science and clinical researchers who utilize diverse models.

## RESULTS

### Demonstration of distinct optical emission paths

Multiparametric imaging depends upon negligible cross-talk between probes, since interference between two fluorescent dyes can lead to erroneous calculations from the acquired signals. Lack of optical crosstalk along the light path was evaluated using a blanking plate, which was used to block either the short/585nm light path or long/710nm light path (Figure 2D). To test for potential dye crosstalk, experiments were performed in which Langendorff-perfused rat hearts were loaded with a single fluorescent probe (Rhod2-AM <u>*or*</u> RH237), and the degree of cross-talk between the two optical paths was assessed independently (Figure 2E, F). Representative optical signals recorded after staining with Rhod2 are shown in Figure 2E. Note that calcium transients are distinctly visualized with no detectable signal in the 710 nm long-pass channel (RH237 channel). Conversely, staining with RH237 only resulted in robust voltage signals with negligible signal in the 585/40 nm channel (Rhod2 channel) with only a small max amplitude of <3 counts (Figure 2F). There was no detected fluorescence from Rhod2 on the 710 nm long pass channel (0 ± 0 counts) compared to the maximum of 17 ± 1 counts on the 585 nm centered channel (p < 0.0001). With RH237 staining only, there was an average maximum of 37 ± 5 counts compared to the 2 ± 1 counts on the 710 nm longpass and 585 nm centered band, respectively, p < 0.0001). These tests demonstrate that RH237 and Rhod2 signals selectively correspond to transmembrane voltage and intracellular calcium, respectively, with minimal cross-talk between the two probes. The latter is comparable to previously reported dual-sensor configurations (5,13).

### Use of ryanodine receptor antagonist to demonstrate signal specificity

Ryanodine, a ryanodine receptor antagonist, is known to significantly impact intracellular calcium transients (36) triggered by action potentials, with minimal effect on action potential morphology (13). To further demonstrate signal specificity, a subset of experiments were performed on juvenile rats to illustrate the effect of ryanodine on calcium transient versus action potential characteristics (Figure 3). As anticipated, ryanodine-supplementation resulted in a marked reduction in the calcium transient peak amplitude by 60% (Figure 3A,C) with no effect on action potential amplitude (Figure 3B,D). Also expected (37), ryanodine-treatment lengthened both the calcium transient duration by 42% (CaD80, Figure 3G) and the action potential duration by 40% (APD80, Figure 3H). This study further demonstrates the specificity of our optical configuration, and that the acquired RH237 and Rhod2 signals are accurately separated.

**Figure 3:**
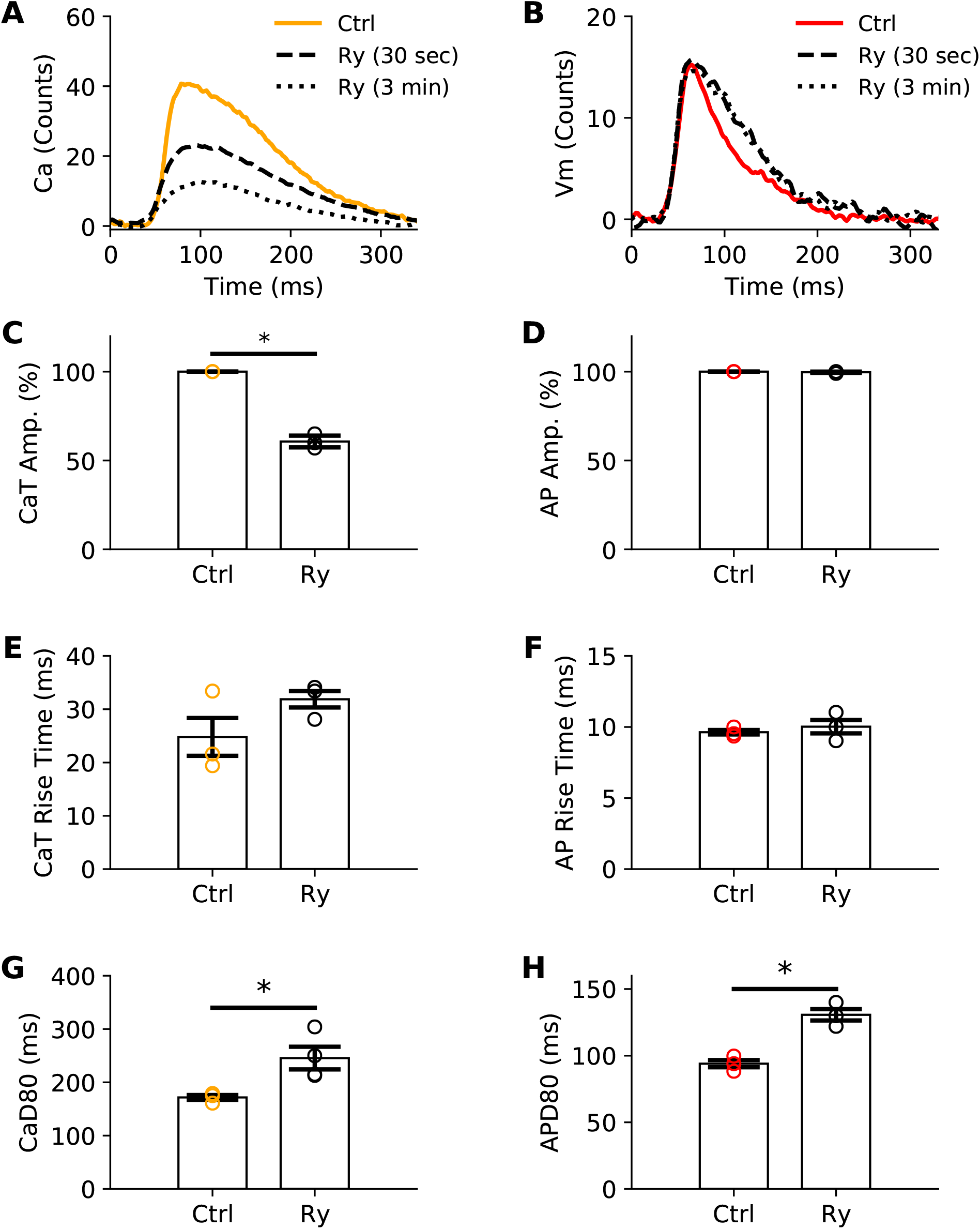
Use of ryanodine receptor antagonist to demonstrate signal specificity. Representative optical action potential **(A)** and calcium transient **(B)** recorded under control perfusion (Ctrl) or immediately after administration of ryanodine (Ry, 30 sec – 3 min). Ryanodine (Ry) administration significantly reduced peak calcium transient amplitudes (**A, C**), with no effect on optical action potential amplitudes (**B, D**). Normalized fluorescent signals were used for time measurements. Calcium transient (**C, E, G**) and action potential (**D, F, H**) parameters are shown before and 1 minute after ryanodine administration at a pacing frequency of 180 msec cycle length. Mean ± SEM, n=3 juvenile rat hearts.

### Simultaneous optical mapping of transmembrane voltage and intracellular calcium

Experimental studies were performed to demonstrate the spatiotemporal performance of our optical mapping system, with suitable signal-to-noise ratio (SNR) for simultaneous dual mapping. Langendorff-perfused rat (2-3 cm length, laying down) or piglet hearts (5-8 cm length, suspended) were loaded with both fluorescent probes (RH237, Rhod2) and optical signals were acquired concurrently (Figure 4). Dye loading can be quickly verified by measuring regions on either image without de-interlacing, which is necessary with other single sensor systems. Following image acquisition (1msec exposure time, 1000 fps, 768×208), optical signals were spatially averaged using a pixel radius of 30 for SNR measurements. These acquisition settings resulted in a SNR of 74 for Rhod2 and 39 for RH237. A slight increase in the exposure time (1.2 msec, 814 fps, 768×256) improved the SNRs to 85 for Rhod2 and 47 for RH237 (see Supplemental Table 2 for SNR calculations). In comparison, when the exposure time was doubled (2 msec, 500 fps, 768×384), the SNR improved to 121 for Rhod2 and 58 for RH237. No difference in fluorescence signal quality was observed within the short image acquisition time (2 sec). Repetitive imaging over the course of 1hr reduced the voltage peak fractional fluorescence (ΔF(p)/F_0_) from 3% to 1.9%, and the calcium peak fractional fluorescence from 4% to 3% (data not shown, n=3).

**Figure 4:**
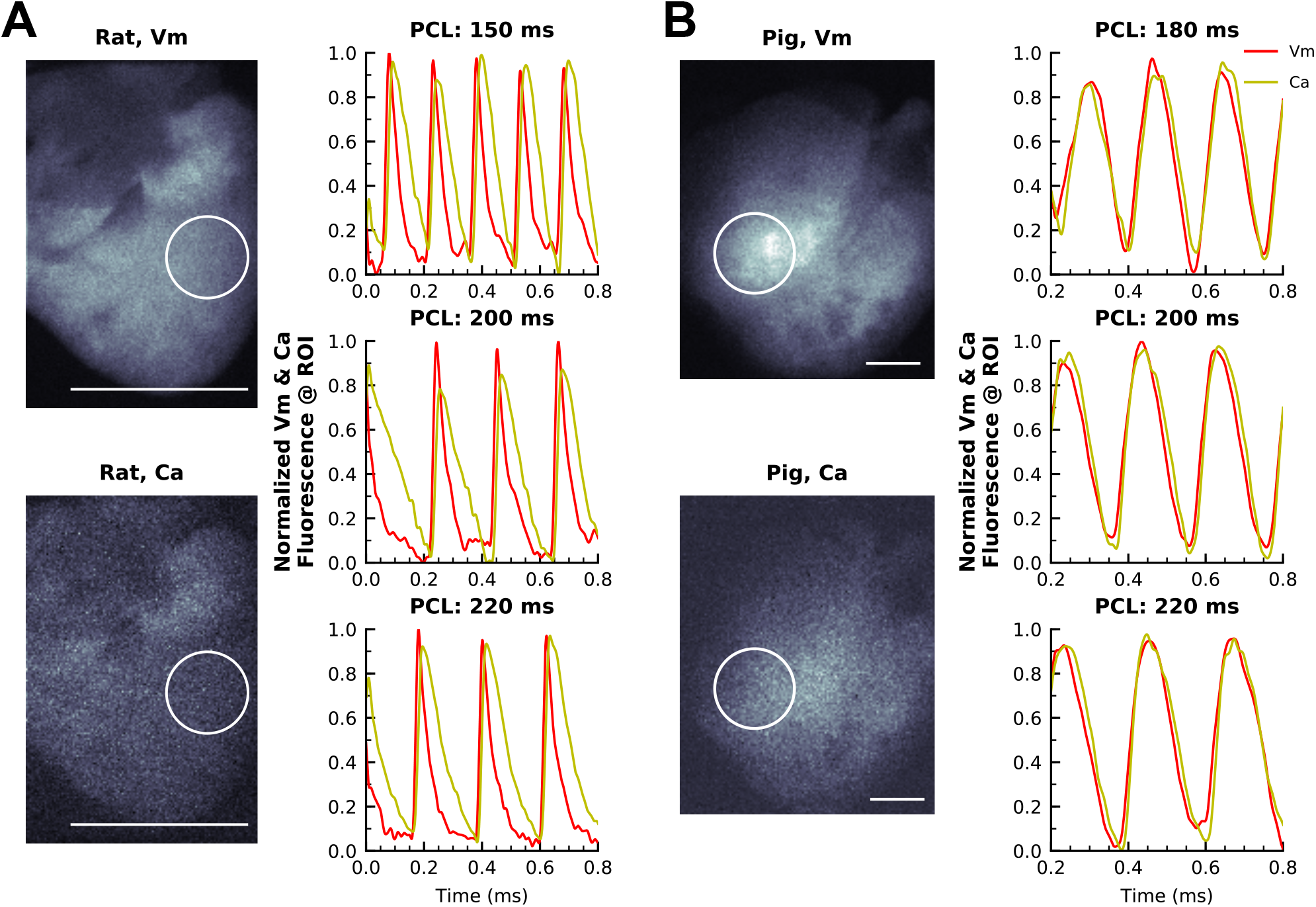
Multiparametric optical signals acquired from Langendorff-perfused rat and piglet hearts. **(A)** Normalized transmembrane voltage (RH237, red) and intracellular calcium (Rhod2, orange) fluorescence signals acquired simultaneously from an excised rat heart during electrical pacing (150, 200, 220 msec pacing cycle length). In this example, each individual image represents 384×256 pixels acquired at 814fps. **(B)** Normalized fluorescence signals acquired simultaneously from excised piglet hearts during electrical pacing (180, 200, 220 msec pacing cycle length). In this specific example, each individual image represents 640×512 pixels acquired at 406fps. Circle denotes region of interest (rat: 45 pixel region, piglet: 80 pixel region in this example). Note: Schneider 17mm f/0.95 lens used for rat hearts, Fujinon 6mm f/1.2 lens used for piglet hearts. Vm = transmembrane voltage, Ca = intracellular calcium signal. ROI = region of interest. Scale bar = 1 cm.

Examples of voltage and calcium signals acquired simultaneously from an isolated rodent heart are shown in Figure 4A, and in Supplemental video 1 & 2. As expected, action potential activation preceded calcium transients for all pacing rates (150, 200, 220 msec cycle length). To demonstrate the versatility of our optical mapping system, supplementary studies were performed on isolated piglet hearts that were imaged in an upright and suspended orientation (180, 200, 220 msec cycle length, Figure 4B, Supplemental video 3 & 4). These experimental studies highlight the utility of our setup for simultaneous transmembrane voltage and intracellular calcium recordings of cardiac preparations, with excellent SNR and temporal resolution.

### Requisite spatiotemporal resolution for whole heart optical mapping

To demonstrate spatiotemporal resolution of our optical mapping system, we assessed excitation-contraction coupling and measured epicardial conduction velocity. Calcium and voltage images were briefly processed by convolution with a uniform kernel for box blurring of increasing size (Figure 5A-H), as previously described(38). The blurring process decreased salt and pepper noise and improved the overall quality of isochronal maps and videos with minimal effect on spatial resolution (Figure 5I). Box blurred images were used for subsequent image analysis, such as conduction velocity measurements (Figure 6A-D). Calcium and voltage signals were acquired with high temporal resolution upstrokes (Figure 6C), and as expected, isochrones show an elliptical pattern of action potential activation and calcium release that originated at the pacing site (Figure 6A, B). A shorter time delay was observed between voltage and calcium signals at the apex of the heart compared with the base (Figure 6C) - an anatomical difference in excitation-contraction coupling that has previously been described (13). Notably, apex-to-base differences can result in slower conduction velocity measurements when calcium measurements are used as a surrogate for voltage wavefront velocity in isolated whole heart preparations (Figure 6D), as compared to cell monolayer preparations (11,39).

**Figure 5:**
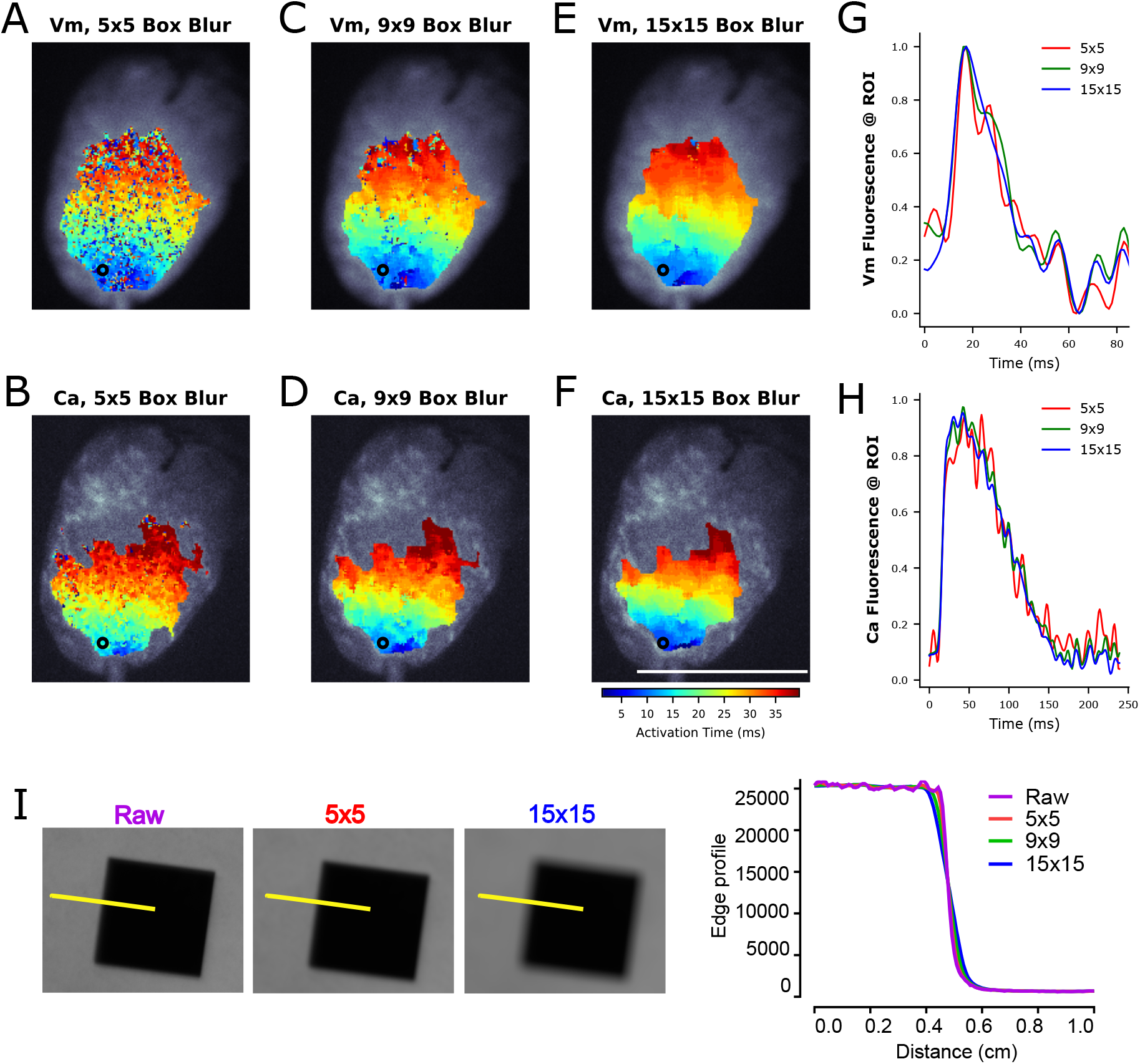
Spatial filtering of optical signals using box blurring. The fluorescence intensity of each pixel is averaged with neighboring pixels. **(A-F)** 384×256 pixel images were convolved with an odd uniform kernel for box blurring to improve SNR for isochrone analysis. Salt and pepper noise is decreased with increasing kernel size for both Vm (top) and Ca (bottom) signals. (**G, H)** Traces show increased SNR with increasing kernel size. Note: The displayed signal is measured from a single pixel (ROI = region of interest), whereas typically for signal analysis an averaged region of interest is used consisting of several pixels from images that have not been box blurred. **(I)** Spatial resolution is demonstrated in the raw image (purple), 5×5 box blurred image (red) and 15×15 box blurred image (blue). Corresponding edge profile trace illustrates spatial resolution of approximately 0.8mm (raw, 5×5 box blurring), 0.86mm (9×9 box blurring) to 1.2mm (15×15 box blurring). Scale bar = 1cm

**Figure 6:**
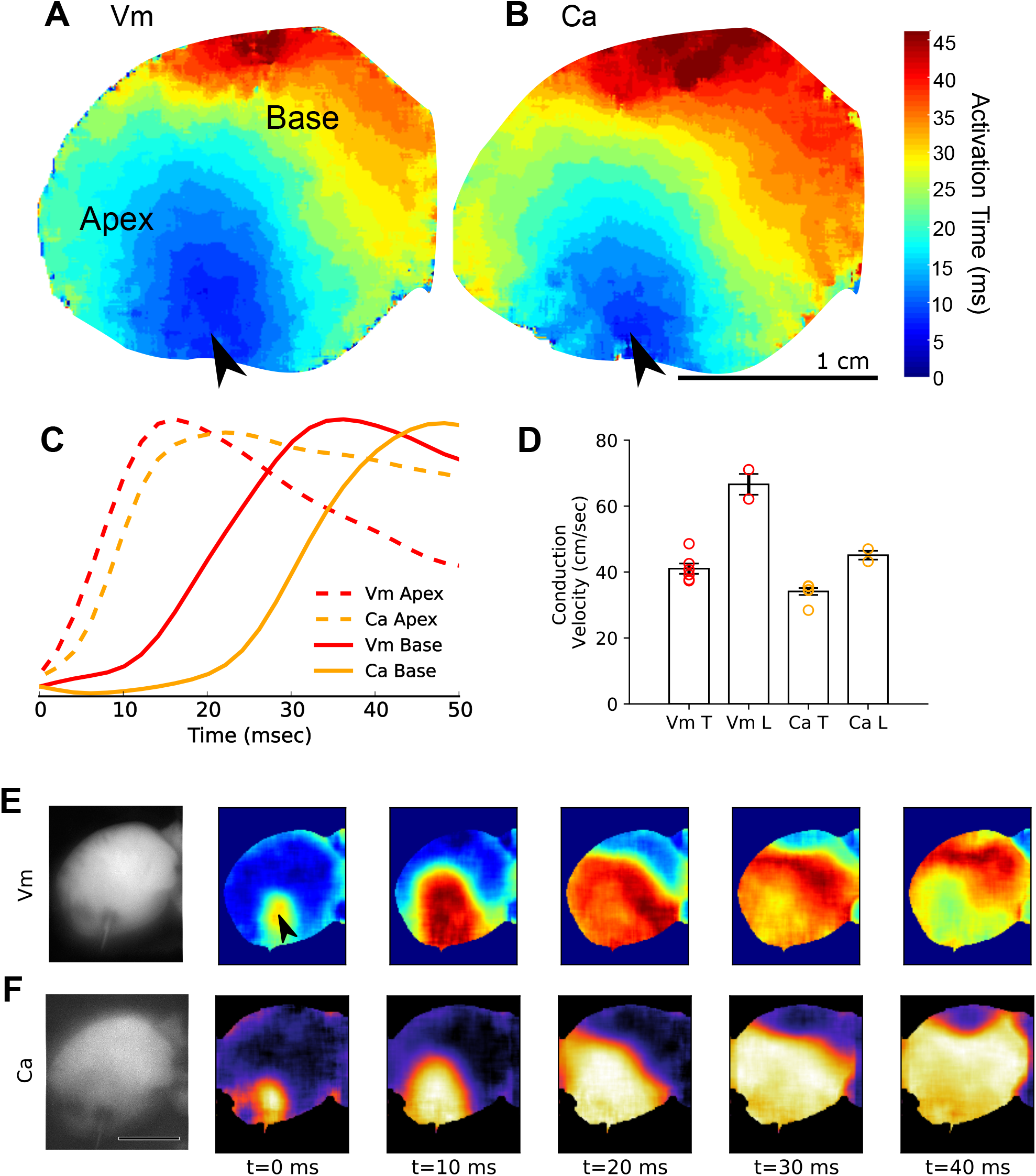
Images from simultaneous acquisition of voltage and calcium signals. (**A**) Voltage activation isochronal map. (**B**) Calcium release isochronal map. (**C**) Voltage and calcium upstrokes display anatomical differences in excitation-contraction coupling (apex versus base). (**D**) Transverse (T) and longitudinal (L) conduction velocity computed from voltage and calcium wavefront propagation. Longitudinal propagation velocity of the calcium wavefront was significantly slower than the voltage conduction velocity (t-test, p<0.001). Slower calcium conduction velocities coincide with increased lag time between the action potential and calcium transient near the base of the heart. Conduction velocity was calculated from 6 angles for transverse and 2 angles for longitudinal measurements. (**E**) Sequential images of impulse propagation show the voltage wavefront originating from the pacing electrode (arrow) and propagating toward the base. Spatial hetereogeneity was observed in the activation and repolarization pattern at t = 40 msec (**B**) Sequential images of the calcium wavefront follows the voltage wavefront, with an example of folding observed at t = 40 msec. Arrows denote pacing electrode; Vm = transmembrane voltage, Ca = intracellular calcium signal. Scale bar = 1cm

As an example, an isolated rat heart preparation is shown wherein longitudinal voltage conduction velocity was 67 ± 4.6 cm/sec compared to the 45 ± 2.0 cm/sec for calcium (Figure 6). Transverse voltage and calcium velocities were 41 ± 2.3 and 34 ± 1.6 cm/sec. In this example, spatial heterogeneity was readily observed as the anisotropic ratio (longitudinal/transverse) was 1.62 for voltage conduction velocity, compared to 1.32 for calcium velocity, which illustrates discontinuity in wavefront propagation (Figure 6E,F). Indeed, viewing the time course of the voltage and calcium activation patterns of this heart pinpointed the activity in left, basal region. At t=40 msec, an area showed a discontinuity activation/repolarization pattern (Figure 6E), with a wavebreak clearly visible on the calcium channel (Figure 6F). This anatomical region was consistent with the increased lag of the calcium release following depolarization, and may suggest underlying myocardial damage. Indeed, the described imaging platform provides sufficient spatial resolution to visualize the spread of electrical activity and calcium cycling in cardiac preparations.

### Induction of cardiac alternans to demonstrate spatial resolution

Arrhythmias are relatively uncommon in the rodent heart under basal conditions, due to species differences in cardiac size and ion channel expression (40). Therefore, a burst pacing protocol was employed to induce ventricular tachyarrhythmias for subsequent pathophysiological imaging. Following burst pacing, concordant alternans were readily observed, wherein an alternating rhythm was observed in the action potential duration and/or intracellular calcium concentration (41–44) (data not shown). Alternans can be detected as T-wave alternans on an electrocardiogram (Figure 7A), or identified optically, from voltage and/or calcium signals acquired from a single region of interest (Figure 7B-G). Conversely, discordant alternans arise when alternating rhythms in different spatial regions are out of phase. Such heterogeneities can be identified with optical mapping of fluorescence signals from multiple sites across a cardiac preparation with high spatiotemporal resolution (45). One such example is shown in Figure 7, in which ventricular tachycardia was encountered concomitantly with spatially discordant calcium transient alternans following dynamic pacing (S1-S1, 70 msec cycle length, isolated rat heart). With higher spatiotemporal resolution than previously reported (Table 1), different tissue regions can be identified and investigated using the described single-sensor setup.

**Figure 7:**
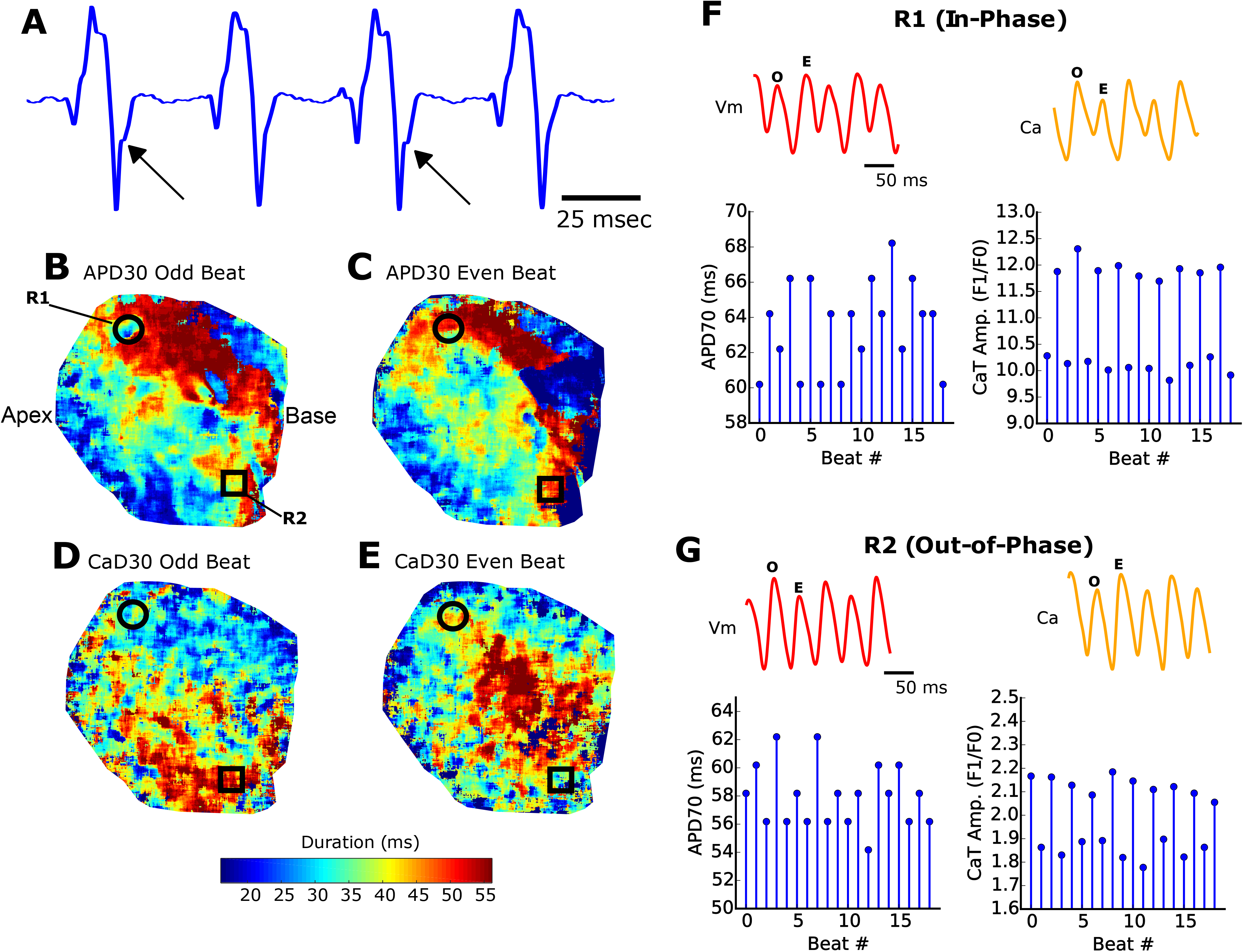
Simultaneous calcium and voltage imaging during ventricular tachycardia. (**A)** T-wave alternans (arrows) on alternating waveforms recording from an isolated, intact heart. **(B, C**) APD30 map shows repolarization heterogeneity during an odd “O” and even “E” beats. (**D,E**) CaD30 signals also show reuptake heterogeneity across the epicardial surface during odd and even beats. Optical action potentials and calcium transients display alternans that can be in-phase **(F)** or out-of-phase **(G)**. In-phase alternans are represented by a small calcium transient amplitude following a shortened action potential duration. Out-of-phase alternans describe a tall calcium transient amplitude that follows a shortened action potential duration. APD30 = Action potential duration at 30% repolarization; CaD30 = Calcium transient duration at 30% relaxation; R1 & R2 = regions of interest; Vm = transmembrane voltage; Ca = intracellular calcium signal. Scale bar = 1cm

Studies suggest that calcium alternans are mechanistically responsible for action potential duration alternans (46), however the association has not been fully elucidated. Depending on the underlying condition and cellular function, two different variants of alternans can manifest. “Positive coupling” describes the scenario in which a small calcium transient amplitude occurs after a shortened action potential (42). Conversely, “negative coupling” describes a tall calcium transient amplitude that is associated with a shortened action potential. In the current study, we show examples of both positive and negative electromechanical coupling (in- and out-of-phase, respectively, Figure 7E,F). We report a novel finding that calcium transient alternans can be spatially discordant, while electrical alternans remain concordant. The described system excels in providing both a wide field of view and high spatial resolution, which facilitated the detection of discordant alternans. Importantly, discordant alternans are considered a precursor for reentry arrhythmias (41), as repolarization gradients become amplified when neighboring cells are out of phase (41,43).

## DISCUSSION

In the current study, we present a novel and innovative approach to dual mapping of cardiac tissue that employs recent advancements in imaging technology. Key advantages include the simplicity of construction (path splitter, single sensor, single light), the elimination of artifacts common for dual light sources and dual camera designs, use of optically compatible dyes, and reduced cost barrier. This dual mapping platform does not require the coordination of exposure frame rate and LED light triggering, which can be technically challenging and also divides the acquisition frame rate by the number of probes (e.g. dual mapping calcium and voltage would otherwise decrease the effective frame rate by 50%). With this configuration, individual images of calcium or voltage are each acquired, simultaneously, at 384×256 resolution at 814 fps. If necessary for a specific application, the acquisition rate can be increased with vertical cropping or the field of view can be expanded to decrease the frame rate, all while maintaining SNR for the detection of calcium and voltage signals.

We validated the spatial and temporal performance of our optical mapping system using isolated whole rat and piglet hearts. Proof-of-principle experiments were performed to measure optical action potentials and calcium transients, spatial distribution of excitation-contraction coupling, conduction velocity measurements in transverse/longitudinal directions, and spatiotemporal resolution of electrical and mechanical alternans. Importantly, accurate separation of transmembrane voltage and calcium signals was shown with single-dye loading experiments and also by application of ryanodine. In the latter, ryanodine administration decreased calcium transient amplitude – with no discernible effect on action potential amplitude (13). We also demonstrated that a larger heart size (piglet, 5-8 cm length) could be accommodated without sacrificing acquisition speed by using wider strips (1024×256) and/or utilizing a wide-angle lens. Notably, the described system has a small footprint that expands its versatility of use between cardiac preparations of different size and orientation. The system configuration can also be outfitted with other camera models to achieve specific temporal and spatial resolution needs. As an example, the N256 camera (SciMedia, Costa Mesa, CA) has a 256×256 sensor at a frame rate of 1818 fps, which could result in dual 256×128 images using the described configuration. Finally, the described approach reduces the investment for a panoramic imaging setup (47–49) by decreasing the total number of required cameras.

One limitation of our approach is the sensitivity of the front-illuminated sCMOS sensor, which peaks at 82% quantum efficiency, but has a roll-off at 700 nm, in the emission spectrum of RH237. We see this limitation as an acceptable trade-off compared to the increased speed that the camera provides compared to back-illuminated options. We did not characterize the loss in quantum yield from the optical path splitter, though we anticipate losses that are comparable to other dichroic based emission splitting systems. The optical path was optimized by using high transmission band (>95%) filters and a high-speed lens (f/0.95 or f/1.4). Another potential limitation of the camera is the small pixel size of 6.5 µm, which is not necessarily required for applications at the tissue level, such as whole heart preparations. We have mitigated the small pixel size by employing box blur algorithms during post-acquisition processing when necessary to improve signal fidelity (binning could also be used). A single excitation light source, single camera platform can complicate the control of dual emission intensities; although we did not encounter this problem due to the similar SNR of the dyes and wide (16-bit) dynamic range of the camera. Finally, a slight difference in the focal plane between 585nm versus 710nm can occur due to chromatic focal shift – although this did not impact fluorescence signals at the tissue level. Color-corrected lenses can be employed to minimize chromatic aberration.

## CONCLUSION

We developed a novel platform to simultaneously acquire voltage and calcium signals from intact, whole heart preparations using an optical path splitter, single camera and single excitation light. Notably, the described platform is composed entirely of off-the-shelf components, which can aid in the adoption and successful implementation of this setup by other laboratories. The described protocol also employs a readily available dye combination (RH237, Rhod-2AM), thereby negating the need for genetically-encoded indicators or fluorescent probes that are not commercially available. We confirm the specificity of our emission signals with negligible cross-talk and demonstrate high spatiotemporal resolution for excitation-contraction coupling, conduction velocity, and concordant/discordant alternans research. The described platform is easy to align and has a small footprint, which provides flexibility in its use for tissue preparations of varying size and/or orientation. In summary, the described platform requires minimal technical complexity, yet provides the spatiotemporal resolution and signal-to-noise ratio necessary for the investigation of voltage/calcium kinetics, conduction velocity and arrhythmia incidence.

## METHODS

Animal procedures were approved by the Institutional Animal Care and use Committee of the Children’s Research Institute, and followed the National Institutes of Health’s <u>*Guide for the Care and Use of Laboratory Animals*</u>. All animals were euthanized by exsanguination under anesthesia during heart excision, as detailed below.

### Isolated rodent heart preparation

Unless otherwise noted, experiments were conducted using adult male Sprague-Dawley rats (2-3 months of age, 250-350g, n=11, Taconic Biosciences). Animals were anesthetized with 3-5% isoflurane, the heart was excised and then transferred to a temperature-controlled (37°C) constant-pressure (70 mmHg) Langendorff perfusion system. Excised hearts were perfused with Krebs-Henseleit buffer throughout the duration of the experiment, as previously described (<1 hour)(12,50,51). Three electrodes were positioned to acquire a far-field electrocardiogram in the lead II configuration. In a subset of experiments, male juvenile rats (postnatal day 5, 10 grams, n=6) were used to demonstrate the effects of ryanodine-supplementation on electromechanical coupling.

### Isolated piglet heart preparation

Yorkshire piglets (2-4 weeks of age, n=4) were used in a supplementary study to demonstrate the versatility of the optical setup. Briefly, an intravenous bolus injection of fentanyl (50μg/kg) and rocuronium (1mg/kg) was administered, and anesthesia was maintained with isoflurane (1-3%), fentanyl (10-25μg/kg) and pancuronium (1mg/kg). The heart was excised and submerged in cardioplegia (4°C) and then flushed with Krebs-Henseleit solution by aortic cannulation. The heart was transferred (~10 minutes at room temperature) to a temperature-controlled (37°C) constant-pressure (70 mmHg) Langendorff-perfusion system. To avoid ischemic-injury in these larger tissue preparations, the heart was suspended in contrast to the “laying” rat heart. Accordingly, the dual mapping platform (camera, path splitter) was relocated in proximity to the larger capacity perfusion system.

Once a baseline heart rate was established (10 min), the perfusate was supplemented with 10 μM (-/-) blebbistatin (Sigma-Aldrich) to reduce motion artifact for subsequent imaging experiments (52,53). Fluorescent dyes were added sequentially, as a bolus, through a bubble trap located proximal to the aortic cannula. Based on predetermined myocardial staining time for each dye, 50 μg Rhod-2AM (1,50) was added first and allowed to stabilize for 10 min, followed by 62.1 μg RH-237 staining for 1 min (12). The myocardial tissue was re-stained by RH237 if needed throughout the duration of the experiment (54). Homogenous dye loading is shown in Supplemental Figure 1. For electrical stimulation, a coaxial stimulation electrode was positioned on the ventricular epicardium, which was driven by a Bloom Classic electrophysiology stimulator (Fischer Medical). Pacing current was set to 1.5x the threshold (resulting in typically 1.8 mA), with 1 msec monophasic pulses.

### Ryanodine administration

To demonstrate negligible cross-talk between voltage and calcium signals, a subset of experiments was performed in the presence of ryanodine (n=3). Ryanodine has previously been shown to selectively affect calcium transient upstroke and amplitude(13). 10 µM ryanodine was added as a bolus, through a bubble trap located proximal to the aortic cannula. Hearts were subjected to continuous pacing during imaging (180 msec), and images were captured before and immediately after application of ryanodine (30 seconds, 1 and 3 minutes).

### Instrumentation

The overall system configuration is shown in Figure 1. The epicardium was illuminated using broad light emitting diode (LED) spotlights centered at 530 nm (Mightex), equipped with an optical filter to constrict the excitation band (ET530/40x nm, Chroma Technologies). LED radiant power varied between 50 – 200 mW, as needed. At the onset of imaging, the excitation LEDs were automatically enabled via a direct TTL connection from the camera output. Due to the use of a single light source for excitation, complex light triggering was not needed to coordinate alternating LED light sources with individual frames.

An image splitting device (OptoSplit II, Cairn Research, Ltd) was positioned in front of a sCMOS camera (Andor Technology, PLC, Zyla 4.2 PLUS); the corresponding light path is shown in Figure 1B. The beam splitter was configured with a dichroic mirror (660+nm, Chroma Technologies, see Supplemental Table 1) that passed RH237 emission and reflected Rhod2 fluorescence. High transmission emission filters were used for Rhod2 (ET585/40m, Chroma Technologies) and RH237 emitted light (long pass ET710, Chroma Technologies). A fixed focal length 17mm/F0.95 lens was attached to the front of the imaging splitting device (Schneider, #21-010456) for rat hearts, and a wide-angle 6mm/F1.2 lens (Fujinon, #DF6HA-1B) was used for pig hearts. Details of the optical configuration are shown in Figure 1B,C and the experimental workflow is displayed in Figure 1D.

*MetaMorph* v7.10.2.240 (Molecular Devices, LLC) was used for camera configuration, optosplit image alignment and LED on/off triggering. To guide manual optosplit alignment, *MetaMorph* overlays the live images as contrasting colors or as subtractive greyscales to highlight misalignment. With this live feedback, images are quickly aligned (<1 minute) using the optosplit’s “long” and “short” control knobs (Figure 2). As an alternative to *MetaMorph*, images could also be aligned with the free software *μManager* (55). After alignment, any standard image acquisition software can be used such as *MetaMorph, μManager*, or *Solis* (Andor Technologies, software supplied with camera). The acquired image will include two fields, which can be separated using *MetaMorph*, *μManager*, or alternative imaging software that includes automated tools. LEDs were attached to a controller (SLC-SA04-U/S, Mightex) that was triggered “on” 1 second prior to image acquisition. The computer consisted of a Xeon CPU E3-1245 v5 3.50 GHz (Intel corporation), 32 GB of RAM, and a non-volatile memory express solid state disk (NVMe SSD, Samsung 960 Pro). A frame grabber was used for imaging control and acquisition (Karbon #KBN-PCE-CL4-F, BitFlow). Because of the high data rate of acquisition (due to high spatial and temporal resolution and bit depth), the NVMe SSD disk was essential for reducing data rate bottlenecks. The frame grabber with 10-tap CameraLink™ connection was necessary to achieve the fastest frame rates possible; a USB connection would result in much slower frame rates.

To maximize spatiotemporal resolution, the sCMOS camera sensor was cropped and set to an acquisition rate of 814 frames per second (fps). The exposure time to achieve this frame rate was 1.206 msec, with a 98.4% duty cycle. This configuration resulted in two images from the splitting device, each 384×256 pixels, with a 16 pixel boundary between each image to negate optical crosstalk (Figure 2D). The field of view was approximately 2.1 x 1.4 cm, which was sufficient to fully image pediatric rat hearts (0.3 g average weight, 1.7 cm length from aortic root to apex). For the larger adult rat hearts (1 g average weight, 2.7 cm in length), the field of view was extended by increasing the working distance. The actual pixel size on the sensor is 6.5 µm with a projected pixel size typically 45-80 µm – depending on working distance and field of view from a given lens. The front of the image splitter is a standard C-mount, which allows a user to select any standard lens to use in combination with accessories (back extension tubes, diopters) to adjust field of view and working distance. The projected pixel size was measured for each study and individually calibrated for conduction velocity measurements. If needed, the frame rate could be increased further by vertical cropping; the feasibility of which was tested with 208 vertical pixels at 1000 fps, which resulted in adequate SNR on the epicardial surface (see results section for details). For supplemental piglet heart studies (80-120 g average weight, 5-8 cm length), a wide-angle lens was used in conjunction with a larger area of the sensor to attain a field of view approximately 5.9 × 4.7 cm.

### Signal and image processing

Following image acquisition, signal or image processing was performed as outlined in Figure 1D. Signal processing and data analysis were performed using a custom MATLAB script(26,51). A circular region of interest was taken in the center of the raw image with a 30-pixel radius (5-20 mm^2^ area on the heart, depending on lens), averaged, and plotted against time. Drift removal and temporal smoothing were applied when necessary. Drift removal was performed by subtraction of a polynomial fit (0^th^, 1^st^, or 2^nd^ order). To remove high frequency noise, a 5^th^ order Butterworth low-pass filter was applied to the resulting signals with a cut-off frequency adjusted between 50-150 Hz. After pre-processing, a peak detector was used to measure the total number of action potentials or calcium transients in the file across time. Characteristics from each event are measured and averaged, including action potential duration, calcium transient duration, time to 90% peak, and amplitude as described previously(26,50,56,57).

Image processing was performed and isochrone maps were constructed in the *rhythm* software(38). The background was removed, convolved with a uniform kernel for box blurring(38) and then time signals were low-pass filtered below 100 Hz. A similar approach was previously described by Laughner, et al. 2012 in which a 3×3 kernel size was used to process images acquired from a MiCAM ULTIMA (100×100 pixels, 100 μm pixel size). We chose a larger uniform kernel size (15×15) since our camera sensor (Zyla 4.2) has sharper resolution (1024×1024 pixels, 6.5 μm pixel size). The activation time of every pixel on the heart was defined as the maximum derivative of the action potential or transient upstroke, which was plotted for both voltage and calcium, respectively. As proof of concept, we also tested the feasibility of measuring conduction velocity across the epicardial surface by exporting the activation maps and analyzing via ORCA(58). Subsequent electrical wave propagation images were constructed using custom Python scripts and plotted with matplotlib(59). Signal-to-noise ratio (SNR) of the time-series was calculated as the action potential amplitude (ΔF) over the standard deviation of the baseline during the diastolic interval(60). The number of pixels used to calculate the SNR was noted when necessary; importantly, the SNR can be improved considerably after box blurring and/or spatial averaging.

### Statistical analysis

Statistical tests were performed using the R software package. Datasets from the ryanodine experiments were compared pairwise using Welch’s T-test and significance reported as p < 0.05.

## Supporting information

Supplemental Video 1 - Rat Voltage

Supplemental Video 2 - Rat Calcium

Supplemental Video 3 - Piglet voltage

Supplemental Video 4 - Piglet calcium

## ABBREVIATIONS

APD_30_: action potential duration at 30% repolarization
Ca: intracellular calcium signal
CaD_30_: calcium transient duration at 30% relaxation
MEHP: mono-2-ethylhexyl phthalate
Rr: ryanodine
SNR: signal to noise ratio
Vm: transmembrane voltage

## DECLARATIONS

### Ethics approval and consent to participate

Animal procedures were approved by the Institutional Animal Care and use Committee of the Children’s Research Institute, and followed the National Institutes of Health’s <u>*Guide for the Care and Use of Laboratory Animals*</u>.

### Consent for publication

Not applicable

### Availability of data and material

The datasets used and/or analyzed during the current study available from the corresponding author on reasonable request.

### Competing Interests

The authors declare that they have no competing interests.

### Funding

This work was supported by the National Institutes of Health (R00ES023477 to N.G.P, R01HL139472 to N.G.P), Children’s Research Institute and Children’s National Heart Institute. We thank the generosity of the NVIDIA corporation for the graphics processing unit to perform CUDA-enabled image processing. Funding sources had no role in the design of the study, collection, analysis, interpretation of the data or writing the manuscript.

### Author contributions

RJ, DM, BS, LS, JH, DM, NGP performed experiments; RJ, DM, BS, NGP analyzed data; RJ, DM, NGP prepared figures; RJ and NGP drafted manuscript, RJ and NGP conceived and designed experiments; RJ, DM, BS; LS, JH, DM and NGP approved manuscript.

## Acknowledgements

The authors gratefully acknowledge Narine Sarvazyan, Charles Berul, and Yu-Ling Shao for helpful discussions, Manelle Ramadan and Morgan Burke for technical assistance, and Nobuyuki Ishibashi and Takuya Maeda for biological materials.

## SUPPLEMENTAL FILE LIST

### Supplemental Video 1

#### Transmembrane voltage mapping of rat heart epicardium

An adult rat heart was imaged during ventricular pacing at cycle length 250 msec. The voltage wavefront can be seen originating from the center of the posterior ventricle and propagating across the surface, followed by a slight delay and retrograde atrial conduction. Calcium imaging was performed simultaneously (see Supplement Video 2). The images were box blurred using a 15 x 15 uniform kernel and the length of the heart is approximately 2.5 cm from base to apex.

### Supplemental Video 2

#### Simultaneous calcium mapping of rat heart epicardium

The calcium activity of the adult rat heart was mapped concurrently with transmembrane voltage. The calcium wavefront can be seen following the voltage propagation from Supplement Video 1. The images were box blurred using a 15 x 15 uniform kernel and the length of the heart is approximately 2.5 cm from base to apex.

### Supplemental Video 3

#### Transmembrane voltage mapping of pig heart epicardium

A juvenile pig heart was imaged during an episode of ventricular tachycardia (cycle length = 150 msec). Circus movement with wave collision can be observed. Calcium imaging was performed simultaneously (see Supplement Video 4). The images were box blurred using a 15 x 15 uniform kernel and the length of the heart is approximately 5.0 cm from base to apex.

### Supplemental Video 4

#### Simultaneous calcium mapping of pig heart epicardium

A juvenile pig heart was imaged during an episode of ventricular tachycardia (cycle length = 150 msec). Circus movement with wave collision can be observed. Transmembrane voltage imaging was performed simultaneously (see Supplement Video 3). The images were box blurred using a 15 x 15 uniform kernel and the length of the heart is approximately 5.0 cm from base to apex.

## Notes

#### Summary of Updates

Text and title modifications

