## Supplemental Materials for "Lights, Camera, Path Splitter: A New Approach for Truly Simultaneous Dual Optical Mapping of the Heart with a Single Camera"

**Running title:** Dual optical mapping

**Keywords:**

Optical mapping, calcium cycling, transmembrane voltage, electrophysiology

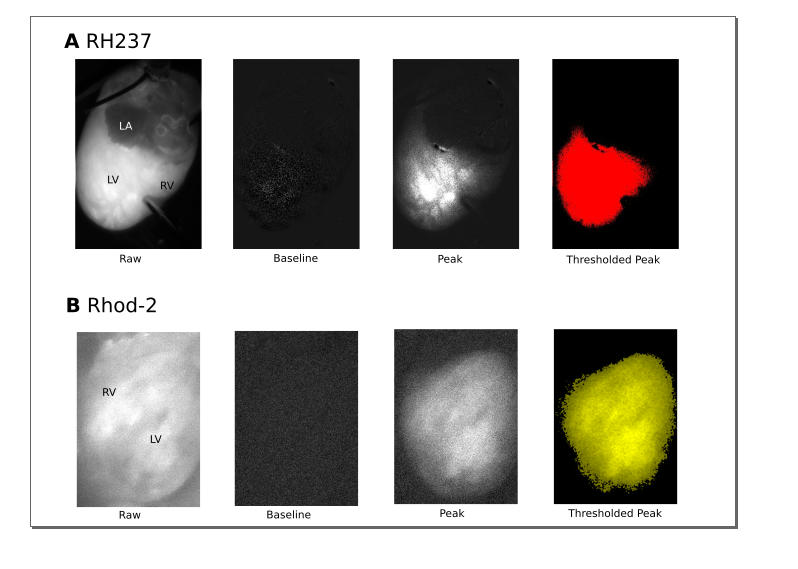

**Supplemental Figure 1.**  **Single dye loading homogeneity.** Fractional fluorescence was calculated by dividing each fluorescent image in the series by the average baseline image (ΔF/F_0_). The resultant fractional fluorescence images show near-black during baseline and bright during peak activity (peak of the action potential or calcium transient). The lookup table was not modified between baseline and peak images. A threshold was performed to show highest area of staining in pseudo-color. (A) RH237 was independently loaded to the heart. (B) Rhod-2 was independently loaded to a different heart to ensure specificity of the dye-loading. Examples of both anterior and posterior orientation are shown. LA = left atrium, LV = left ventricle, RV = right ventricle.

**Supplemental Table 1**: Major components, specifications, part numbers, and manufacturers for dual mapping system

| **Description** | **Specs (Part #)** | **Manufacturer** |
| --- | --- | --- |
| Excitation (2x) | 530 nm ± 20 nm (ET530/40x) | Chroma Technology (Bellows Falls, VT) |
| Emission Dichroic | 660+ nm (T660lpxrxt) | Chroma Technology (Bellows Falls, VT) |
| Rhod-2 Emission | 585 nm ± 20 nm (ET585/40m) | Chroma Technology (Bellows Falls, VT) |
| RH237 Emissions | 710+ nm (ET710lp) | Chroma Technology (Bellows Falls, VT) |
| sCMOS Camera | Full-frame 100 fps (Zyla 4.2 PLUS) | Andor Technology (Belfast, Ireland) |
| Excitation LEDs | 530 nm peak, 200 mW (PLS-0530-030-15-S) | Mightex Systems (Toronto, CA) |
| Lens | 17mm, f/0.95, (21-010456) | Schneider Optics (Hauppauge, NY) |
| Lens | 6 mm, f/1.2 (DF6HA-1B) | Fujifilm (Tokyo, Japan) |
| Image Splitter | Two channel (OptoSplit II) | Cairn Research Ltd (Kent, UK) |

**Supplemental Table 2**: Quantified performance by signal-to-noise ratio measurements with different exposure times and image processing.

| **Exposure Time, msec (FPS)** | **Image Processing** | **Rhod-2AM (Ca)** | **RH237 (Vm)** |
| --- | --- | --- | --- |
| 1.0 (1000) | 30 px radius mean | 74 | 39 |
| 1.2 (814) | 30 px radius mean | 85 | 47 |
| 2.0 (500) | 30 px radius mean | 121 | 58 |
| 1.2 (814) | 15 x 15 box blur | 2.3 | 12.1 |
| 1.2 (814) | 15 x 15 box blur with 100 Hz LPF | 3.9 | 14.7 |

**Supplemental Video 1**

**Transmembrane voltage mapping of rat heart epicardium.** An adult rat heart was imaged during ventricular pacing at cycle length 250 msec. The voltage wavefront can be seen originating from the center of the posterior ventricle and propagating across the surface, followed by a slight delay and retrograde atrial conduction. Calcium imaging was performed simultaneously (see Supplement Video 2). The images were box blurred using a 15 x 15 uniform kernel and the length of the heart is approximately 2.5 cm from base to apex.

**Supplemental Video 2**

**Simultaneous calcium mapping of rat heart epicardium.** The calcium activity of the adult rat heart was mapped concurrently with transmembrane voltage. The calcium wavefront can be seen following the voltage propagation from Supplement Video 1. The images were box blurred using a 15 x 15 uniform kernel and the length of the heart is approximately 2.5 cm from base to apex.

**Supplemental Video 3**

**Transmembrane voltage mapping of pig heart epicardium.**  A juvenile pig heart was imaged during an episode of ventricular tachycardia (cycle length = 150 msec). Circus movement with wave collision can be observed. Calcium imaging was performed simultaneously (see Supplement Video 4). The images were box blurred using a 15 x 15 uniform kernel and the length of the heart is approximately 5.0 cm from base to apex.

**Supplemental Video 4**

**Simultaneous calcium mapping of pig heart epicardium.** A juvenile pig heart was imaged during an episode of ventricular tachycardia (cycle length = 150 msec). Circus movement with wave collision can be observed. Transmembrane voltage imaging was performed simultaneously (see Supplement Video 3). The images were box blurred using a 15 x 15 uniform kernel and the length of the heart is approximately 5.0 cm from base to apex.
